# The tyrosine phosphatase PRL regulates attachment of *Toxoplasma gondii* to host cells and is essential for virulence

**DOI:** 10.1101/2022.01.24.477641

**Authors:** Chunlin Yang, William J. Blakely, Gustavo Arrizabalaga

**Affiliations:** Department of Pharmacology and Toxicology, Indiana University School of Medicine, Indianapolis, Indiana, USA; Department of Microbiology and Immunology, Indiana University School of Medicine, Indianapolis, Indiana, USA

**Keywords:** PRL, Toxoplasma, tyrosine phosphatase, virulence, attachment

## Abstract

The pathogenesis of *Toxoplasma gondii* is mainly due to tissue damage caused by the repeating lytic cycles of the parasite. Many proteins localized to the pellicle of the parasite, particularly kinases, have been identified as critical regulators of the *Toxoplasma* lytic cycle. However, little is known about the associated protein phosphatases. Phosphatase of regenerating liver (PRL), a highly conserved tyrosine phosphatase, is an oncogene that plays pivotal roles in mammalian cells and typically associates with membranes via a conserved prenylation site. PRL in *Toxoplasma* has a predicted prenylation motif in the C-terminus like other homologs. We have determined that TgPRL localizes to the plasma membrane and that disruption of TgPRL results in a defect in the parasite’s ability to attach to host cells. This function is dependent both on TgPRL’s membrane localization and phosphatase activity. Importantly, in vivo experiments have shown that while mice infected with parental strain parasites die within days of infection, those infected with parasites lacking TgPRL not only survive but also develop immunity that confers protection against subsequent infection with wild-type parasites. Immunoprecipitation experiments revealed that the PRL-CNNM (Cyclin M) complex, which regulates intracellular Mg^2+^ homeostasis in mammalian cells, is also present in *Toxoplasma*. Consistent with this interaction, parasites lacking TgPRL had higher intracellular Mg^2+^ levels than the parental or complemented strains, suggesting TgPRL is involved in regulating intracellular Mg^2+^ homeostasis. Thus, TgPRL is a vital regulator of the *Toxoplasma* lytic cycle and virulence, showing its potential as a target of therapeutic intervention.

**IMPORTANCE:** Infection with *Toxoplasma gondii* can lead to severe and even life-threatening diseases in people with compromised or suppressed immune systems. Unfortunately, drugs to combat the parasite are limited, highly toxic, and ineffective against the chronic stage of the parasite. Consequently, there is a strong demand for the discovery of new treatments. A comprehensive understanding of how the parasite propagates in the host cells and which proteins contribute to the parasite’s virulence will facilitate the discovery of new drug targets. Our study meets this objective and adds new insights to understanding the lytic cycle regulation and virulence of *Toxoplasma* by determining that the protein phosphatase TgPRL plays a vital role in the parasite’s ability to attach to host cells and that it is essential for parasite virulence.

## INTRODUCTION

*Toxoplasma gondii* is a single-celled parasite of the phylum Apicomplexa, capable of infecting any warm-blooded animal, including approximately 30% of humans worldwide (1, 2). Humans are infected congenitally or by ingestion of either environmental oocysts, shed in the feces of cats, or tissue cysts in undercooked meat of infected animals. Most infections are asymptomatic during the acute stage, but to evade the immune response, the parasite converts to a latent encysted form, thus establishing a chronic infection (3). In immunocompromised individuals, such as lymphoma and AIDS patients, new infections or reactivation of parasites in pre-existing cysts can lead to severe toxoplasmic encephalitis (4). In addition, for congenital infections, given the immature nature of the fetal immune system, toxoplasmosis can lead to blindness, severe neurological problems, and even death (5, 6).

A significant portion of the pathogenesis associated with toxoplasmosis is a direct consequence of the repeating cycles of invasion, division, and egress that drive the propagation of the parasite (7, 8). Both invasion and egress of the parasite are active events that rely on regulated secretion from specialized organelles and on the parasite’s gliding motility system (9, 10). Secretion and motility are tightly regulated by opposing effects of cGMP and cAMP and by calcium signaling (11–14). Particularly, calcium-dependent phosphorylation plays a key role in the regulation of the parasite’s lytic cycle (15–18). While many of the kinases involved in regulating the effectors of the lytic cycle have been elucidated (19–21), little is known about the role of phosphatases. Strong candidate phosphatases for roles in lytic cycle regulation are those that associate with the parasite’s pellicle and cytoskeleton as they are critical for both motility and secretion.

Recently we characterized serine/threonine protein phosphatases predicted to be membrane-associated and determined that only PPM5C, a PP2C family protein phosphatase, localizes to the plasma membrane, where it regulates attachment of *Toxoplasma* to host cells (22).

One member of the tyrosine protein phosphatases of *Toxoplasma* that may also be associated with the pellicle is a homolog of the phosphatase of regenerating liver (PRL). PRL is a highly conserved tyrosine phosphatase that possesses a prenylation motif at the C-terminus (23). Typically, prenylation and the polybasic region preceding it guide PRL homologs to the plasma membrane (23). PRL was first identified in regenerating livers because of its highly elevated expression (24), hence its name. Mammalian cells encode three PRLs, which show significant amino acid sequence identity. N-terminal epitope-tagged mouse PRLs were shown to be present in the inner leaflet of the plasma membrane and in intracellular punctate structures that were determined to be early endosomes (25). However, studies using antibodies against mammalian PRLs have identified variable localizations. For instance, PRL1 was found to be present in the endoplasmic reticulum and mitotic spindle (26), and PRL3 and PRL1 were found in the plasma membrane and cytoplasm (27). Multiple studies have indicated that PRLs are oncogenic and are overexpressed in a wide variety of cancer cells, especially in metastatic lesions (28–30). As their expression levels in cancer cells closely correlate with disease progression, PRLs are considered potential therapeutic targets for cancer treatment (31). Although PRLs play vital roles in the regulation of cancer progression, the molecular mechanisms behind their function remain unclear (23, 31). Studies have shown PRL is involved in a breadth of cell signaling pathways, including Rho-family GTPase, PI3K-Akt, Ras-MAPK, STATs, P53, and Src/ERK1/2 (23, 31). However, no clear substrates of PRL in these signaling pathways have been identified so far.

Despite the important role of PRL in mammalian cells, its function in apicomplexan parasites is poorly understood. The *Plasmodium falciparum* homolog PfPRL is 218 amino acids in length and contains the conserved C-terminal signature CAAX motif “CHFM” for prenylation. In vitro phosphatase activity assays indicated that PfPRL is a tyrosine phosphatase, as it is preferentially inhibited by a tyrosine phosphatase inhibitor (32). Immunofluorescence assay using antibodies against PfPRL showed that PfPRL is present in the endoplasmic reticulum and a subcompartment of the food vacuole when the parasite is in intraerythrocytic stages (32). By contrast, the *Toxoplasma* homolog (TgPRL) has not been characterized. TgPRL is slightly larger in size than the human and *Plasmodium* homologs with 404 residues and has a C-terminal prenylation motif “CAIM”. In this study, we determined that the TgPRL associates with the parasite membrane and that it regulates attachment to host cells. Importantly, deletion of *TgPRL* renders the parasite avirulent, which reveals the potential of this protein as a target for therapeutic intervention.

## RESULTS

### TgPRL is localized to the plasma membrane

Like its homologs in human cells, TgPRL has a C-terminal prenylation motif, indicating that it is likely membrane-associated (Fig. 1A). In addition, a predicted palmitoylation site of four amino acids upstream of the prenylation motif increases the likelihood of its localization in the plasma membrane (Fig. 1A). To determine the localization of TgPRL, we constructed an ectopic expression plasmid that uses TgPRL’s promoter to drive expression of TgPRL carrying a hemagglutinin (HA) epitope tag at the N-terminus (Fig. 1B). A parasite strain stably carrying this construct (*Δhx:*HA-PRL) was established, and expression of HA-tagged TgPRL at the expected size (45.08 kDa) was confirmed by western blot (Fig. 1C). Immunofluorescence assay (IFA) of the resulting strain showed that the N-terminal HA-tagged TgPRL mainly localizes to the plasma membrane as expected, with some punctate staining in the cytoplasm (Fig. 1C). Interestingly, the N-terminal HA-tagged TgPRL appears to be present only in non-dividing parasites but not in parasites with developing daughter parasites (Fig. 1D).

**Figure 1.**
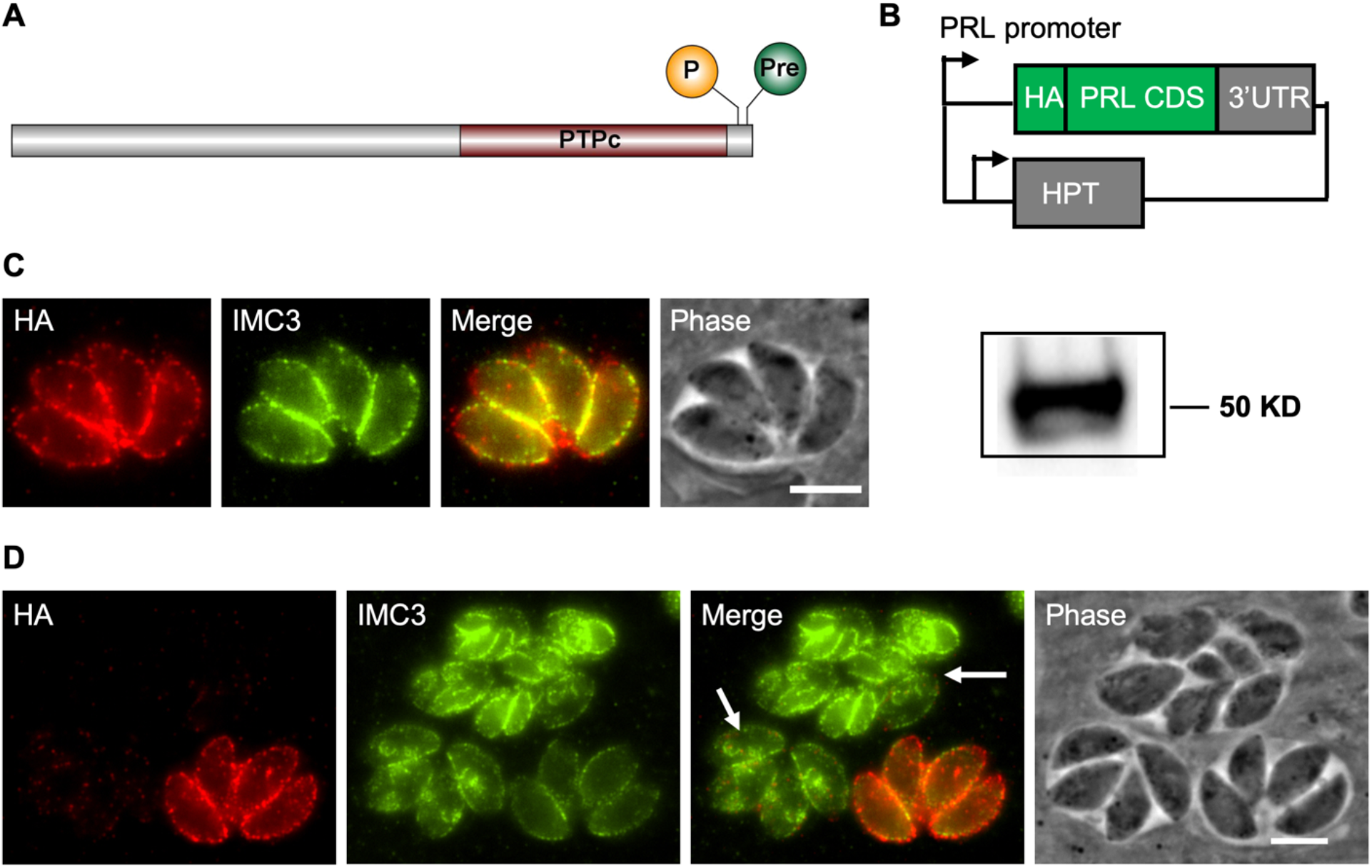
TgPRL localizes to the plasma membrane. (A) Schematic of TgPRL showing the palmitoylation (P) and prenylation (Pre) sites and the tyrosine phosphatase domain (PTPc). (B) Diagram of HA-PRL expression plasmid. One copy of an HA epitope tag is included at the N terminal of the TgPRL coding sequences. Expression of the fused HA-PRL is under the control of the *TgPRL* promoter. The plasmid also contains the selectable marker HXGPRT (HPT). (C) Intracellular parasites expressing HA-PRL were stained with antibodies against the HA epitope (red) and the IMC3 (green). Scale bar = 2 μm. The panel to the right of (C) shows the western blot of parasite extracts probed for HA. (D) Panels show both dividing (arrow) and not dividing HA-PRL expressing parasites stained as in C.

### Knockout of TgPRL affects *Toxoplasma* propagation

To determine the function of TgPRL in *Toxoplasma*, we generated a knockout strain of TgPRL using a CRISPR/Cas9 driven approach (Fig. 2A) (33). Briefly, we constructed a plasmid, pSag1-Cas9-U6-sgPRL, which expresses Cas9 and a guide RNA targeting the first exon of the *TgPRL* gene, and generated a PCR amplicon containing a selectable dihydrofolate reductase (DHFR) marker (34) flanked by overhangs with homology to *TgPRL* sequences upstream and downstream of the Cas9 cut site (Fig. 2A). Both the plasmid and the amplicon were transfected into parasites lacking KU80 (Δ*ku80*) (35) to favor homologous recombination (Fig. 2A). PCR with primers flanking the cut site was performed, and the resulting amplicons were sequenced to validate the successful insertion of the selection marker (Fig. 2B). To attribute any defect observed in the Δ*prl* parasite strain, we introduced an N-terminal HA-tagged TgPRL at the 5’UTR of the disrupted *KU80* gene for complementation (Fig. 2C). The resulting complemented strain Δ*prl*.*cp* was confirmed for the expression of HA-PRL by both western blotting (Fig. 2D) and IFA (Fig. 2E).

**Figure 2.**
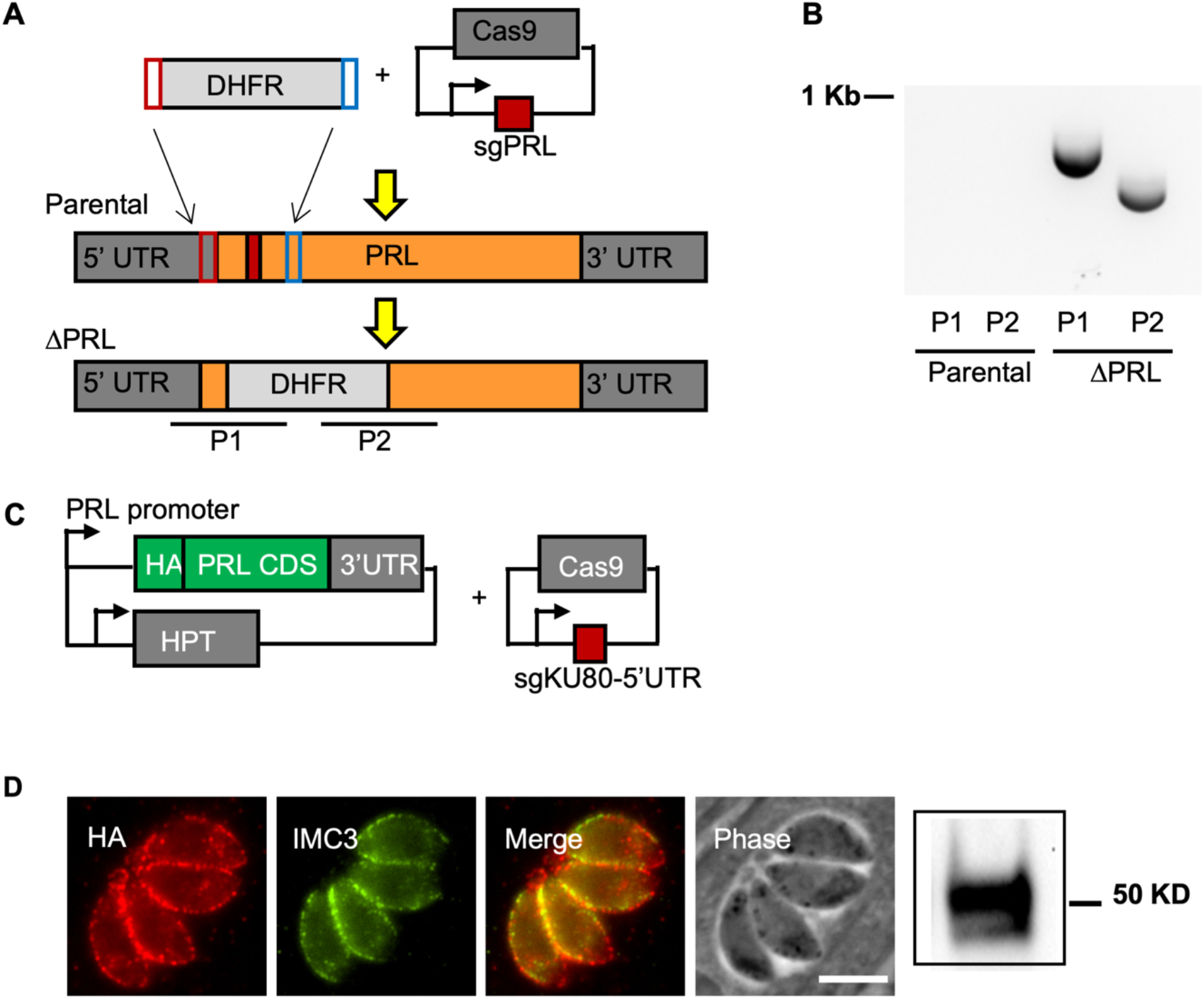
Generation and complementation of TgPRL knockout strain. (A) Diagram depicts the strategy used for developing the TgPRL knockout strain Δ*prl*. Parasites were transfected with a plasmid encoding Cas9 and a guide RNA that targets the first exon of *TgPRL* (red box) and an amplicon that contains the selectable marker DHFR and regions of homology to the *TgPRL* locus. The bottom graphic represents the resulting edited genome in the knockout strain. P1 and P2 are the regions amplified by PCR to confirm disruption. (B) Results of PCR of both parental and Δ*prl* strains for amplicons P1 and P2 are shown. (C) Diagram depicts the plasmids used to develop TgPRL complementation strain Δ*prl*.*cp*. The HA+PRL expression plasmid is the same one shown in Figure 1B. A plasmid that expresses Cas9 and a guide RNA targeting the 5’UTR of KU80 gene was used to locate the expression cassette. (D, E) Expression of HA+PRL in the complemented strain was confirmed by immunofluorescence assays (E) and western blot (D).

To examine the effect of disrupting TgPRL on parasite propagation, we performed plaque assays of the parental (Δ*ku80*), knockout (Δ*prl*), and complemented (Δ*prl*.*cp*) strains. This assay showed that Δ*prl* parasites formed significantly smaller plaques than the parental strain, indicating that disruption of TgPRL affects parasite propagation (Fig. 3A, B). On average, the knockout strain cleared 47.8%±7.6%, while the complemented strain cleared 72.5%±5.7% of the cell monolayer relative to the parental strain. This data is consistent with the predicted fitness score (−2.32) of TgPRL obtained from a genome-wide CRISPR screen (36). Importantly, the expression of an N-terminal HA-tagged TgPRL in the knockout strain was able to partially complement the phenotype confirming the link between the knockout of TgPRL and the phenotype (Fig. 3A, B).

**Figure 3.**
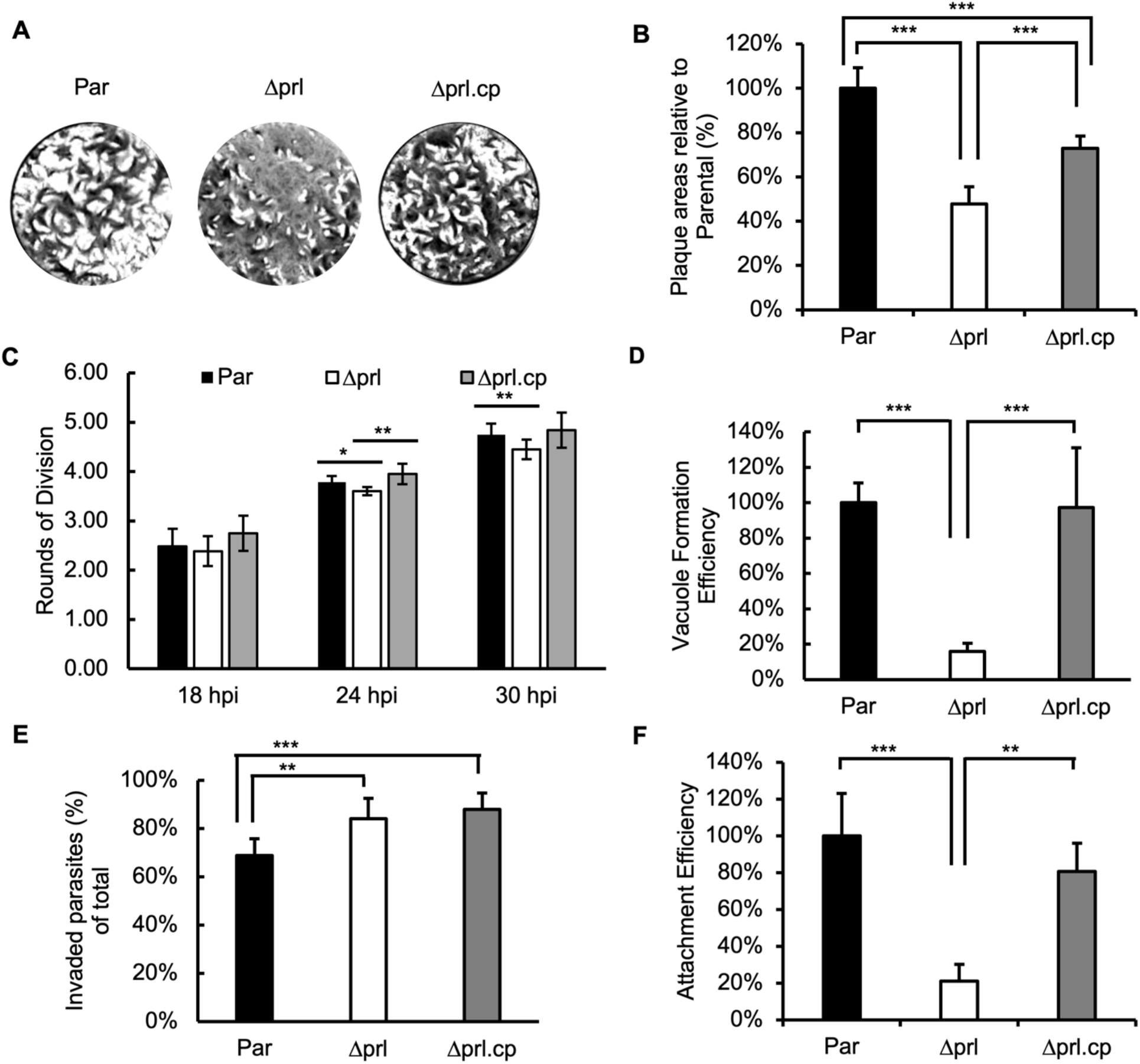
Phenotypic analysis of TgPRL knockout strain. Assays were performed to determine the ability of the knockout (Δ*prl*) strain in propagation, replication, invasion, and attachment parasites with comparison to the parental (Par) and complemented (Δprl.cp) strains. (A) Representative images of plaque assays are shown. (B) The graph represents the quantification of area cleared by plaques relative to parental strain for all strains. (C) For each strain, we quantified the average number of rounds of division at 18-, 24-, and 30-hours post-infection (hpi). (D) Parasites of each strain were allowed to invade cells for 30 minutes, and the number of vacuoles formed was quantitated. Data are shown as % efficiency in relation to the parental strain. (E) Parasites were allowed to invade for 30 minutes, and parasites that were attached but not invaded and those that were invaded were differentially stained as to quantitate invasion. The bars represent the percentage of invaded parasites in the 10 randomly selected fields of view. (F) Results from the same experiment as E were analyzed to quantitate attachment efficiency across all three strains. The bars represent the total number of parasites, including both attached and invaded in the 10 randomly selected fields of view. The data presented are normalized to the average of the data obtained from parental parasites. For all data graphs, n = 3 biological replicates x 3 experimental replicates and p-value was estimated by two-tailed Student’s t-test. Error bars show standard deviations (SD). * p-value < 0.01; ** p-value < 0.001; *** p-value < 0.0001.

### Knockout of TgPRL causes a defect in host cell attachment

Reduction in plaque size can be due to a disruption in any of the multiple steps that make up the *Toxoplasma* propagation cycle, including attachment, invasion, division, egress, and motility (8). Accordingly, we performed a series of more precise phenotypic assays to identify the specific step affected by *TgPRL* genetic disruption. To compare the division rate of the different strains, we allowed parasites to infect host cells for 30 minutes, after which cultures were incubated for 18, 24, or 30 hours before fixation. For each data point, we scored the number of parasites per vacuole for at least 50 vacuoles and converted the average parasite per vacuole value to the corresponding rounds of cell divisions (Fig. 3C). This assay showed that the Δ*prl* parasites are slightly behind division rounds at 24 hours (Par: 3.78 vs. Δ*prl*: 3.60) and 30 hours (Par: 4.75 vs. Δ*prl*: 4.45) post-infection (Fig. 3C). While small, these differences are statistically significant and complemented by reintroducing the wild-type copy of the gene.

Next, we analyzed the ability of Δ*prl* parasites to enter host cells and establish parasitophorous vacuoles as compared to that of the parental and complemented strains. We found that the number of vacuoles formed by Δ*prl* parasites was 15.9%±4.5%, while the complemented strain formed 97.3%±33.8% of those relative to the parental strain (Fig. 3D). These results suggest that the propagation phenotype observed with the Δ*prl* parasites is likely due to the defect in parasite entry into host cells.

The mutant parasite’s reduced ability to form vacuoles could be due to inefficient attachment, invasion, or both. Accordingly, we performed an invasion assay, in which the outside non-invaded but stably attached parasites were stained green with antibody against the cell surface antigen Sag1 before permeabilization of the cell membrane, and all parasites (both inside and outside) were stained red with anti-Mic5 antibody after membrane permeabilization (37). In this way, we can compare the efficiency of attachment (total number of parasites, both inside and outside) and invasion (the percentage of parasites detected inside the cell, i.e. inside/total) between strains. After this differential staining, we compared the percentages of parasites detected inside cells across the three strains, which showed that the Δ*prl* and complemented parasite strains have a slightly higher percentage of invaded parasites (Fig. 3E). Nonetheless, we noted a significant difference across strains when we quantitated the total number of parasites that either remain outside or are inside the cells. The total number of Δ*prl* parasites detected was 21.0%±9.1% relative to parental, while the complemented strain detected was 80.7%±15.2% relative to the parental (Fig. 3F), which was consistent with the result of the vacuole formation assay above. This assay indicates that disruption of TgPRL severely impairs the parasite’s ability to attach to host cells.

The establishment of a tight attachment of the parasites to host cells is mediated by several adhesion proteins secreted from micronemes in a calcium-dependent manner (38, 39). To determine whether the defect in the attachment of the Δ*prl* parasites is the result of deficient microneme secretion, we monitored secretion of the microneme marker MIC2 across the parental, mutant, and complemented strains. We detected no difference in the constitutive secretion of MIC2 among the three strains (Fig. S1). In addition, we also detected no difference in the amount of calcium-induced secretion of MIC2 among the three strains (Fig. S1). Moreover, there was also no significant difference in secretion from the dense granules, another set of secretory organelles, determined by monitoring Gra1 (Fig. S1). Taken together, these data suggest that the role of TgPRL in attachment is likely independent of microneme secretion.

### Localization and activity are important for TgPRL’s role in attachment

To explore whether the localization of TgPRL in the plasma membrane is necessary for its role in host cell attachment, we mutated the HA-PRL expression plasmid so that the conserved cysteine predicted to be prenylated was mutated to alanine (C401A). The C401A plasmid was transfected into the Δ*prl* parasites using the same approach as for the complementation with the wild-type TgPRL described above. IFA of the established parasite strain showed that the C401A mutant TgPRL was distributed throughout the cytoplasm and no detectable association with the plasma membrane was observed (Fig. 4A), suggesting the prenylation site of TgPRL is required for its localization. Importantly, the mislocalized C401A mutant TgPRL was unable to complement the plaque-forming phenotype of Δ*prl* parasites (Fig. 4B). This indicates that membrane association is critical for TgPRL’s function in the lytic cycle.

**Figure 4.**
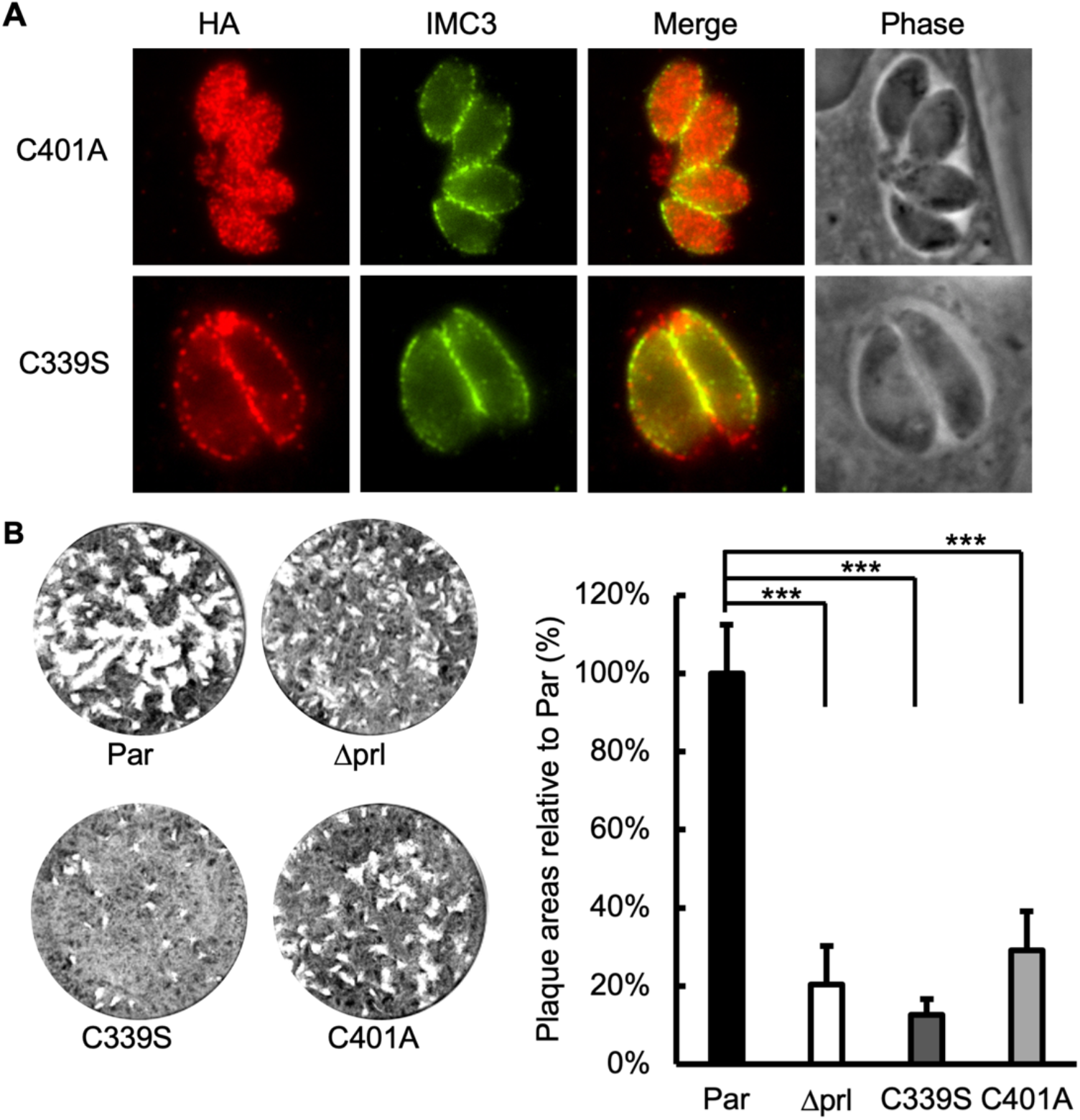
Role of membrane localization and phosphatase activity in TgPRL function. The knockout strain Δ*prl* was transfected with mutant versions (C339S and C401A) of the complementation plasmid shown in Fig. 2C. (A) Intracellular parasites expressing the C339S or C401A mutant TgPRL were stained to determine the localization of exogenous mutant TgPRL. As expected, C339S correctly associates with the plasma membrane, and C401A is mislocalized to the cytoplasm of the parasite. (B) Representative images of plaque assay of corresponding strains. The chart on the right is the quantification of the plaque area relative to parental. For all data, n = 3 biological replicates x 3 experimental replicates and p-value was estimated by two-tailed Student’s t-test. Error bars show standard deviations (SD). * p-value < 0.01; ** p-value < 0.001; *** p-value < 0.0001.

Typically, a cysteine at the center of tyrosine phosphatase domains is required for establishing a phosphoryl-cysteine intermediate (PTP-Cys-PO_3_) and thus phosphatase function (40). Accordingly, to determine if the phosphatase activity is important for TgPRL’s role in attachment, we mutated the conserved cysteine in the center of the phosphatase domain into serine (C339S) in the HA-PRL expression plasmid and transfected it into the Δ*prl* parasites for complementation. IFA of the resulting strain showed that the C339S mutant TgPRL localized correctly to the plasma membrane (Fig. 4A). Nonetheless, the plaque assays showed that the C339S mutant formed only 12.7%±3.9% plaques relative to the parental, while the Δ*prl* parasites formed 20.4%±9.9%, indicating that the C339S mutant TgPRL not only failed to rescue the phenotype but further decreased plaque-forming deficiency of the mutant strain (Fig. 4B). In conjunction, these results indicate that TgPRL phosphatase activity at the periphery of the parasite is required for the protein’s function during host cell attachment.

### Identification of TgPRL’s potential substrates and interactors

To identify potential substrates and interactors of TgPRL, we performed a co-immunoprecipitation (Co-IP) assay with an HA-PRL expressing strain. Since the interaction between phosphatases and their substrates can be transient and unstable, we used cross-linking with DSSO (disuccinimidyl sulfoxide) prior to immunoprecipitation to stabilize protein-protein interactions. Co-IP was performed by using mouse anti-HA magnetic beads to capture HA-PRL and interacting proteins in the lysate of Δ*prl*.*cp* parasites, with the parental Δ*ku80* parasites as a control. The proteins pulled down by the beads were identified by using mass spectrometry (MS). The obtained data (Dataset 1) was curated with the following criteria to develop a list of putative TgPRL substrates or interactors: no less than five peptides and fold change of peptides over control >=8. Applying these criteria resulted in a list of 18 putative TgPRL interactors (Table 1).

**Table 1.** Putative TgPRL-interacting protein partners and substrates identified by Co-Immunoprecipitation and mass spectrometry.

| ID | Function Annotation | Fold Change | Number of Peptides |  |
| --- | --- | --- | --- | --- |
| | | | Control ($\Delta ku80$) | $\Delta prl.cp$ |
| TGGT1_208718 | putative protein tyrosine phosphatase type IVA A (PRL) | INF | 0 | 54 |
| TGGT1_287980 | FHA domain-containing protein | INF | 0 | 17 |
| TGGT1_202840 | FHA domain-containing protein | INF | 0 | 12 |
| TGGT1_262150 | kelch repeat and K<sup>+</sup> channel tetramerisation domain containing protein | INF | 0 | 11 |
| TGGT1_278660 | putative P-type ATPase4 | INF | 0 | 9 |
| TGGT1_211350 | CBS domain-containing protein (CNNM1) | INF | 0 | 8 |
| TGGT1_297520 | proteophosphoglycan PPG1 | INF | 0 | 8 |
| TGGT1_219700 | putative DNA replication licensing factor MCM4 | INF | 0 | 6 |
| TGGT1_227800 | EF-hand domain-containing protein | INF | 0 | 6 |
| TGGT1_243920 | putative DNA replication licensing factor MCM5 | INF | 0 | 6 |
| TGGT1_263130 | putative citrate synthase | INF | 0 | 6 |
| TGGT1_289970 | hypothetical protein | INF | 0 | 5 |
| TGGT1_311830 | hypothetical protein | INF | 0 | 5 |
| TGGT1_314400 | putative pyruvate dehydrogenase E1 component, beta subunit | 10 | 1 | 10 |
| TGGT1_219860 | putative replication licensing factor | 9 | 1 | 9 |
| TGGT1_227560 | putative IWS1 transcription factor | 9 | 1 | 9 |
| TGGT1_253940 | CAM Kinase family, incomplete catalytic triad | 9 | 1 | 9 |
| TGGT1_305510 | hypothetical protein | 9 | 1 | 9 |
| TGGT1_311625 | WD domain, G-beta repeat-containing protein | 9 | 1 | 9 |

Interestingly, one of the most abundant prey proteins in the list is a CBS domain-containing protein (TGGT1_211350), which contains an unknown function 21 (DUF21) domain near the N-terminus, followed by a cystathionine-β-synthase (CBS) domain. Homology search in the human genome revealed that its homologs are cyclin M (CNNM) family proteins, which are plasma membrane metal transport mediator proteins involved in regulating intracellular Mg^2+^ homeostasis (41). Previous studies have shown that the CNNM family proteins are Mg^2+^ transporters that maintain intracellular Mg^2+^ levels by extruding Mg^2+^ from cells (41). Interestingly, PRLs were identified to interact with cyclin M (CNNM) family proteins physically, and the PRL-CNNM complex plays a significant role in regulating intracellular Mg^2+^ levels (42, 43). The identification of the *Toxoplasma* homolog by TgPRL Co-IP indicates that the PRL-CNNM interaction is conserved in *Toxoplasma*.

### Knockout of TgPRL increases intracellular Mg^2+^ levels

Studies in mammalian cells have shown that the CNNM family proteins maintain intracellular Mg^2+^ levels by extruding Mg^2+^ from cells (41). Through direct interaction with CNNMs, PRL regulates Mg^2+^ levels by inhibiting their Mg^2+^ efflux activity (42, 43). It also showed that PRL overexpression increases intracellular Mg^2+^ levels, and knockdown of PRL decreases Mg^2+^ levels (41). To explore if TgPRL plays a similar role in *Toxoplasma*, we compared intracellular Mg^2+^ levels among the parental, Δ*prl*, and Δ*prl*.*cp* strains by using the magnesium indicator Magnesium Green. Briefly, freshly lysed parasites from each of the three strains were incubated with Magnesium Green for 30 min before washes to remove extracellular Magnesium green. The washed parasites were resuspended with Magnesium Green free medium and incubated for another 30 min to complete de-esterification of intracellular AM esters. Parasites from the three strains were resuspended to the same concentration, and the same volume of parasites was added to a 96-well plate for fluorescence measurements. Surprisingly, the fluorescence signal of Δ*prl* parasites was 38.2% ± 15.6% stronger than that of the parental, while that of the complemented strain was similar to that of the parental, suggesting that disruption of TgPRL increased intracellular Mg^2+^ levels, confirming a role for TgPRL in magnesium homeostasis (Fig. 5).

**Figure 5.**
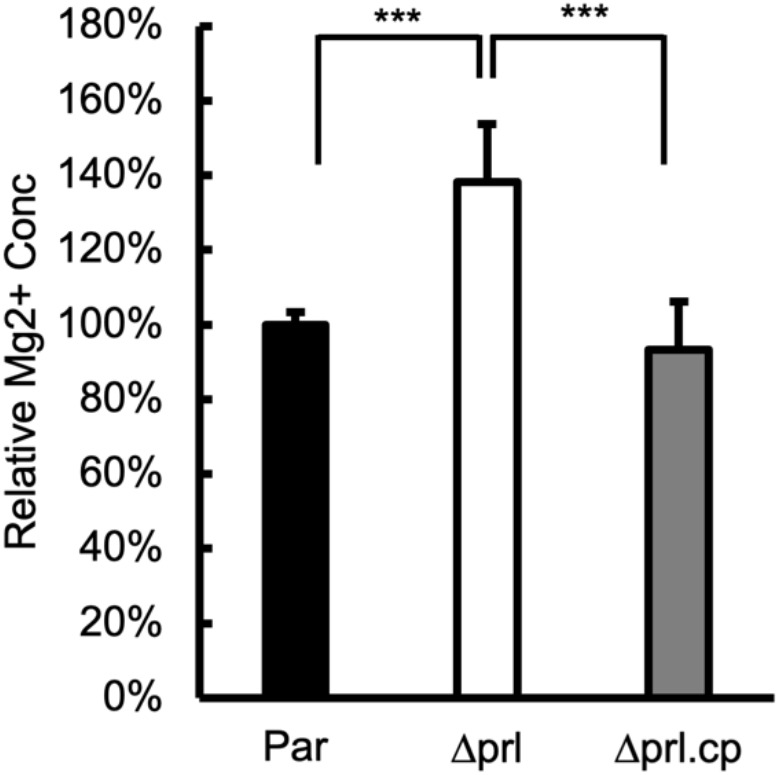
TgPRL knockout parasite strain has a higher intracellular Mg^2+^ concentration. Bars show the relative intracellular free Mg^2+^ concentrations of the parental (Par), knockout (Δ*prl*), and complemented (Δ*prl*.*cp*) strains. The data were from n = 4 biological replicates × 3 experimental replicates. For each biological replicate, data were normalized to the average of the parental. Error bars show standard deviations (SD). p-values are shown between the compared groups. *** indicates p-value < 0.0001.

### Parasites lacking TgPRL is avirulent to mouse

We have found TgPRL is not essential but that its disruption affects parasite propagation in tissue culture, mainly due to a defect in host cell attachment. To explore if the tissue culture propagation phenotype would manifest as an effect on parasite virulence in vivo, we injected CBA/J mice with either parental (Par), Δ*prl* (KO), and Δ*prl*.cp (Comp) parasites to mice. Each parasite strain was injected into two groups of five mice, one with 100 parasites and the other with ten parasites. An extra group of mice was injected with PBS as a control. As expected, all mice infected with the parental strain died, with those injected with 100 parasites succumbing by day eight after injection and those injected with ten by day 9 (Fig. 6). In each of the two groups of mice injected with Δ*prl*.*cp* parasites (Comp 100 and Comp 10), three mice died on day 9 (Fig. 6). Of the four remaining mice, three survived in good health except one in the Comp 10 group, which died on day 20 (Fig. 6). Analysis of sera from the four surviving mice found no antibodies against parasites, suggesting that they were likely not infected (Fig. S2). Remarkably, all mice injected with the Δ*prl* parasites survived, whether they received 100 or 10 parasites (Fig. 6). All but one of the blood samples from the mice injected with the Δ*prl* parasites tested positive for anti-*Toxoplasma* antibodies (Fig. S2), indicating that the mice were efficiently infected but that the Δ*prl* strain is avirulent in mice.

**Figure 6.**
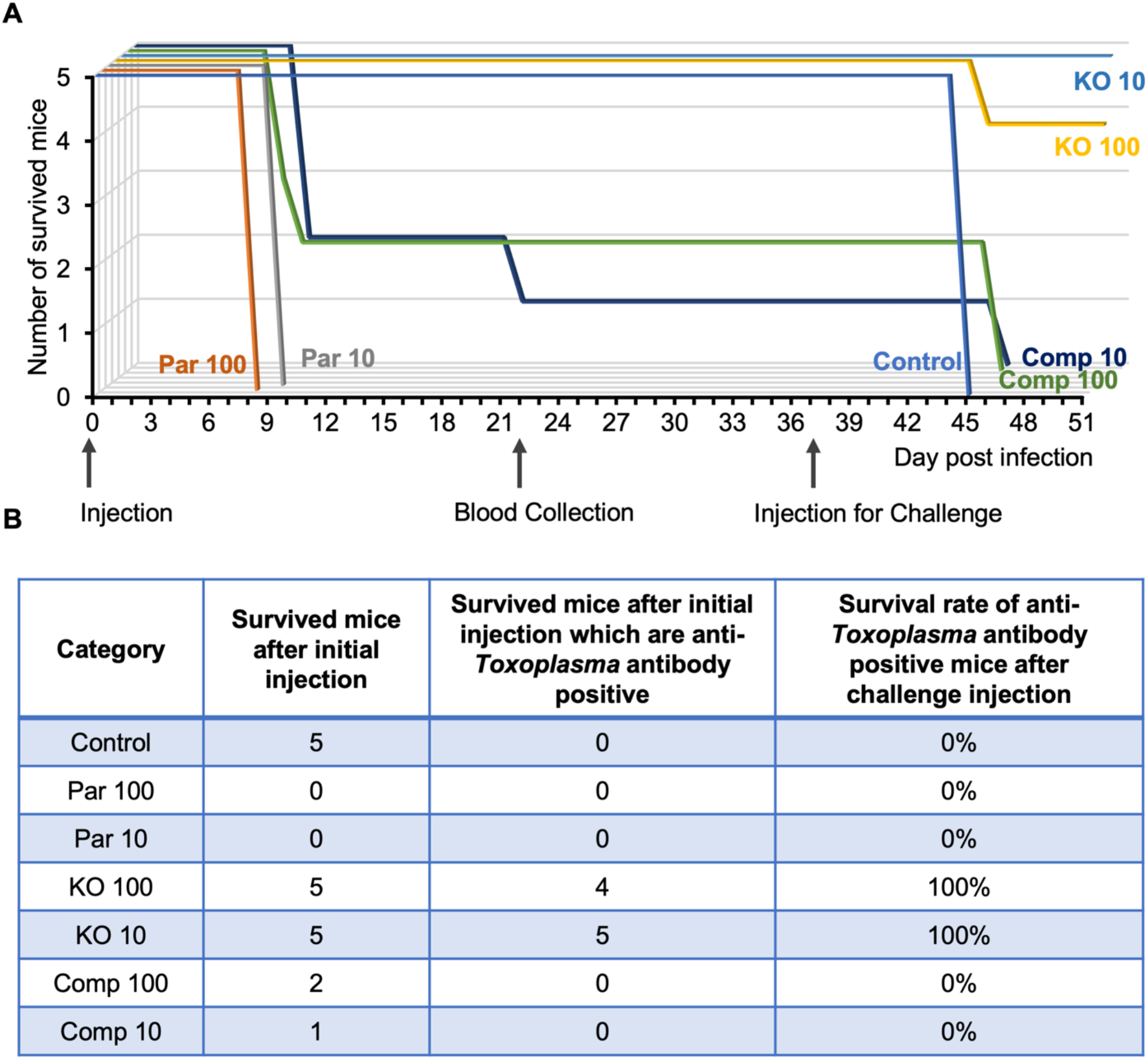
TgPRL knockout strain is avirulent in mice. Mice were separated into groups, and each was injected with either 10 or 100 parasites of the parental (Par 10 and 100), knockout (KO 10 and 100), and complemented (Comp 10 and 100) strains. (A) Animals were monitored after infection, and the day of death was recorded. The graph shows the survival curves of the mice for each of the infected groups. The three arrows pointing at the x-axis indicate the days of the initial injection, blood collection, and challenge injection, respectively. (B) The table shows the number of surviving mice in each test group mice after the initial injection, which tested positive for anti-*Toxoplasma* antibodies, and the survival rate of anti-*Toxoplasma* antibody-positive mice after challenge injection.

To investigate if the antibodies produced by these surviving mice could confer protection against a subsequent *Toxoplasma* infection. We challenged all the surviving mice with an injection of 100 parental parasites two weeks after blood collection. Unsurprisingly, all the antibody-negative mice died on day eight after the challenge. Excitingly, all nine antibody-positive mice that had been infected with the Δ*prl* strain survived in good health (Fig. 6). Taken together, this in vivo experiment showed that disruption of TgPRL renders the parasite avirulent to mice and that infection with Δ*prl* parasites immunizes mice.

## DISCUSSION

In the present study, we characterize the tyrosine phosphatase PRL in *Toxoplasma*. Similar to mammalian homologs, N-terminal epitope-tagged TgPRL localizes mainly to the plasma membrane with some punctate localization in the cytoplasm. Mutation of the cysteine residue in the prenylation motif deprives TgPRL of plasma membrane localization, suggesting that prenylation is required to associate TgPRL to the plasma membrane. Knockout of *TgPRL* is not lethal to the parasites but affects their growth in tissue culture, primarily due to a defect in host cell attachment. The link between TgPRL and host cell attachment adds new insights to our understanding of attachment, one of the key lytic cycle steps of *Toxoplasma* (8). For intracellular parasites, efficient entry of host cells is essential to parasitize a wide range of host cells. *Toxoplasma* can parasitize almost any warm-blooded animal because of its unique gift to attach to and invade host cells. For *Toxoplasma*, adhesion proteins secreted from micronemes help the parasite establish a tight interaction with the host cell (38, 44), and the glideosome centered motility system provides a strong impetus for entering the host cell (45). Recent studies have recognized that *Toxoplasma* attachment to host cells appears to be regulated by more than just microneme secretion. In our previous study, we found that PPM5C, a protein phosphatase of the PP2C family, plays an important role in attachment to host cells that is unrelated to microneme secretion (22). In addition, Calcineurin, a PPP family protein phosphatase, is necessary for both *Toxoplasma* and *Plasmodium falciparum* to strongly attach to host cells before entry, which is also not related to microneme secretion (46). In this study, we have determined that the tyrosine phosphatase TgPRL is required for efficient host cell attachment in *Toxoplasma*. Complete knockout of TgPRL reduces the parasite’s ability to attach firmly to the host cell by almost 80%. However, knockout of TgPRL did not seem to affect microneme secretion. While this result suggests TgPRL’s role in attachment is independent of microneme secretion in general, it is plausible that TgPRL directly or indirectly regulates specific proteins involved in the process. Similarly, other aspects of the parasite’s biology, such as shape, membrane rigidity, and osmotic pressure, could influence attachment and be the target of TgPRL regulation.

Interestingly, we have determined that the disruption of TgPRL increases intracellular free Mg^2+^ concentration in *Toxoplasma*. A role for PRL phosphatases in regulating magnesium homeostasis has been reported in mammalian cells (43). This function appears to be driven by a direct interaction between PRL and cyclin M (CNNM) family proteins (42, 47). CNNM family proteins contain two evolutionarily conserved domains, with a transmembrane unknown function 21 (DUF21) domain at the N-terminus, which spans the plasma membrane, followed by a cystathionine-β-synthase (CBS) domain (41). Previous studies have shown that the CNNM family proteins are Mg^2+^ transporters that maintain intracellular Mg^2+^ levels by extruding Mg^2+^ from the cell and that the formation of PRL-CNNM complex inhibits Mg^2+^ efflux (41). This interaction is dependent on a conserved aspartic acid residue in the CBS domain of CNNM, and mutation of this key amino acid blocks the PRL-CNNM interaction and results in the release of inhibition of Mg^2+^ efflux (43). Moreover, PRL-CNNM interaction seems to be regulated by the phosphorylation status of the cysteine in the center of the phosphatase domain of PRL (43). It showed that phosphorylation of this cysteine blocks the interaction and that phospho-cysteine levels change in response to intracellular Mg^2+^ levels (43).

*Toxoplasma* encodes two proteins with both DUF21 and CBS domains (TGGT1_211350 and TGGT1_307580). Of these two, TGGT1_211350 is most similar to mammalian homologs, hence the name CNNM1. Interestingly, unbiased identification of proteins that interact with TgPRL revealed CNNM1 as one of the top hit proteins, suggesting the PRL-CNNM complex is also present in *Toxoplasma*. If present, this complex may also regulate intracellular Mg^2+^ homeostasis in the plasma membrane, as it does in mammalian cells. Unfortunately, various attempts to add an epitope tag to the C-terminus of endogenous CNNM1 or to express exogenous CNNM1 with epitope tag at the N-terminus or between the DUF21 and CBS domains failed, which hindered our efforts to study this interaction in Toxoplasma further. Nonetheless, our observation that Mg^2+^ levels are disrupted in the TgPRL mutant strongly suggests that the interaction between TgPRL and CNNM1 is present in the parasite. Interestingly, disruption of TgPRL resulted in an increase in intracellular free Mg^2+^ rather than the decrease in intracellular Mg^2+^ levels seen when the mammalian homolog is disrupted (41). Thus, it appears that there might be a different function for the PRL-CNNM interaction in *Toxoplasma*. How the dysregulation of Mg^2+^ homeostasis in the TgPRL mutant connects to the defect in attachment is unclear. Plausibly, an increase of intracellular Mg^2+^ concentration in *Δprl* parasite may trigger some effects, such as it may alter the osmotic pressure of the parasite; or possibly allowing parasites to maintain a higher level of ATP-Mg^2+^, while parasites with additional energy supply may be more inclined to remain free; or may over-activate enzymes leading to complex physiological changes; or trigger disturbances in calcium metabolism, which plays a key role in regulating invasion and egress of *Toxoplasma*.

The Co-IP of TgPRL identified other potential interacting proteins with variable functions, suggesting the involvement of TgPRL in multiple pathways. The top two hits of the Co-IP are two unknown FHA domain-containing proteins (TGGT1_287980 and TGGT1_202840), both of which probably play key roles based on their low fitness scores (−3.15 and −3.54) (36). Notably, three proteins in this list are potentially associated with endocytosis in *Toxoplasma* (48). They are TgK13 (TGGT1_262150), the homolog of *Plasmodium* Kelch13 (K13), TgEps15 (TGGT1_227800) and TgPPG1 (TGGT1_297520). A previous study has reported that K13 plays an important role in *Plasmodium* endocytosis and that Eps15 and PPG1 are two partners of K13 that co-localize with K13 in the hemoglobin-containing endocytic compartment in *Plasmodium* (49). Although little is known about endocytosis in *Toxoplasma*, a large number of proteins required for endocytosis are encoded in its genome, including the three shown here (48). The IFA of N-terminal HA-tagged TgPRL showed that in addition to being localized to the plasma membrane, TgPRL is also present in the cytoplasm in a punctate form. Therefore, those puncta with TgPRL localization may be endocytic compartments in *Toxoplasma*.

One of the most significant findings of our characterization of TgPRL is the discovery that this protein is an essential virulence factor. Mice infected with as many as 100 Δ*prl* parasites survive infection and mount immunity against challenges with parental parasites. This avirulent phenotype is remarkable as the parental strain is highly virulent, with one parasite killing mice within ten days (50), and the mutant can propagate effectively in tissue culture, albeit at a slower rate than the parental strain. This result underscores the critical role of TgPRL in the biology of *Toxoplasma* and reveals this protein as a strong candidate for therapeutic intervention. Future studies on the molecular mechanisms underlying TgPRL function will reveal important information about *Toxoplasma*’s virulence and propagation in infected tissues.

## MATERIALS AND METHODS

### Parasite cultures

All parasites were cultured and maintained in human foreskin fibroblasts (HFF) as previously described (22). The growth medium used was Dulbecco’s Modified Eagle Medium (DMEM) supplemented with 10% fetal bovine serum (FBS), 2mM L-glutamine, and 50 ug/ml penicillin-streptomycin. Parasites used in this study include the strain RH lacking hypoxanthine-xanthine-guanine phosphoribosyl transferase (HXGPRT, RHΔ*hxgprt*, referred to here as Δ*hx*) (51) and RH lacking HXGPRT and Ku80 (RHΔ*ku80*Δ*hxgprt*, referred to as Δ*ku80*) (35). To harvest intracellular parasites, infected cells were scraped off and passed through a 27-gauge needle.

### Generation of HA-PRL expression parasite line

For cloning experiments, genomic DNA was isolated from Δ*ku80* parasites, and cDNA was prepared from total RNA isolated from Δ*ku80* parasites as described before (22). To generate the plasmid for exogenous expression of HA-PRL, the TgPRL coding region was amplified by PCR from cDNA using a forward primer that encodes a hemagglutinin (HA) epitope tag coding sequence in frame with *TgPRL*. All primers used in this study are included in Table S1. To drive expression of the exogenous TgPRL, we amplified a ∼800bp genomic DNA region immediately preceding the start codon of the *TgPRL* gene to include the *TgPR*L promoter and 5’UTR (PRL.pr). The PRL.pr and HA-PRL amplicons were cloned into a plasmid backbone containing a tubulin 3’UTR and an HXGPR selection cassette using In-Fusion HD Cloning to obtain the plasmid pPRL.pr-HA-PRL-Tub-HXGPRT. This plasmid was linearized and transfected into the Δ*hx Toxoplasma* strain by electroporation. Stably transfected parasites were selected for in 50 mg/ml MPA and 50 mg/ml xanthine and clones were established by limiting dilution.

### Generation of TgPRL knockout strain

The CRISPR/Cas9 plasmid pSag1-Cas9-U6-sgPRL-HXG was generated by mutating the sgRNA in the pSag1-Cas9-U6-sgUPRT-HXG plasmid (33) (a gift from Dr. David Sibley) into the sgRNA targeting the first exon of *TgPRL* gene by using Q5 Site-Directed Mutagenesis Kit (NEB). A DNA fragment containing the DHFR selection cassette flanked by short homology to the upstream and downstream of the guide RNA target site was amplified using pJET-DHFR (a gift from Dr. Peter Bradley) as a template. The pSag1-Cas9-U6-sgPRL-HXG plasmid and the PCR amplicon were transfected together into Δ*ku80* parasites. The transfected parasites were treated with 1 mM pyrimethamine for selection, and the parasites were cloned by limiting dilution. Clones were tested by PCR, and the resulting amplicons were sequenced to confirm the expected disruption of *TgPRL*.

### Complementation with TgPRL WT and mutated cDNA

The complementation with wild type (WT) TgPRL used the same plasmid, pPRL.pr-HA-PRL-Tub-HXGPRT, used for generation of the HA-PRL expression line. The complementation constructs with mutant TgPRL were generated by introducing synthesized DNA fragments with either the C339S or C401A mutations by In-Fusion HD Cloning into the complementing plasmid digested with NotI and NcoI or NcoI and AflII. The resulting plasmids were named pPRL.pr-HA-PRL.C339S-Tub-HXGPRT and pPRL.pr-HA-PRL.C401A-Tub-HXGPRT.

The TgPRL WT and mutant complementation plasmids were used as templates to amplify the PRL.pr-HA-PRL-Tub expression cassette plus the HXGPRT selection maker cassette for complementation. To direct the resulting amplicon to insert into the 5’UTR of the disrupted KU80 gene in Δ*prl* parasite strain, a CRISPR/Cas9 plapSag1-Cas9-U6-sgKU80.5’UTR-HXGR-HXG was generated by mutating the sgRNA in the pSag1-Cas9-U6-sgUPRT-HXG plasmid into the sgRNA targeting the 5’UTR of KU80 gene by using the Q5 Site-Directed Mutagenesis Kit. Both the amplicon andpSag1-Cas9-U6-sgKU80.5’UTR-HXGR-HXG plasmid were transfected into Δ*prl* parasite strain. The transfected parasites were selected with 50 mg/ml MPA and 50 mg/ml xanthine in culture and then cloned by limiting dilution.

### Immunofluorescence assays and western blots

Both immunofluorescence assays (IFA) and western blots were performed with previously described protocols (22). For this study primary antibodies used were rabbit anti-HA (Cell signaling Technology) and rat anti-IMC3, both used at 1:1000. For IFA secondary antibodies were Alexa Fluor 594 or Alexa Fluor 488-conjugated goat anti-rabbit and goat anti-rat (Invitrogen), both used at 1:2000. Imaging was performed with a Nikon Eclipse E100080i microscope at 100x magnification. For Western blots we used anti-rabbit or anti-mouse secondary antibodies conjugated to IgG horseradish peroxidase (HRP).

### Phenotypic assays

All the phenotypic assays were performed with standard methods previously described (52). Briefly, for plaque assays, each well of a 12-well plate with confluent HFF monolayers was infected with 500 freshly syringe released parasites. Cultures were allowed to be incubated for six days before being fixed with methanol and stained with Crystal Violet. Imaging was performed using a Protein simple imaging system, and the relative plaque areas were measured by ImageJ and the plugin ColonyArea (53). For doubling assays, 2×10^4^ freshly syringe-released parasites were allowed to invade HFFs for 30 minutes. Cultures were washed with DMEM medium four times to remove free parasites and then incubated for 18, 24, and 30 hours before being fixed with methanol and stained with Hema3 Manual Staining System (Fisher Scientific). For each sample, 50 vacuoles were randomly selected, and the number of parasites per vacuole was recorded and converted to corresponding rounds of division.

To perform vacuole formation assays, 5×10^5^ freshly syringe-released and filter-purified parasites were allowed to invade HFFs for 30 minutes. Unstably-attached and unattached parasites were removed by four washes with DMEM, and the cultures were incubated for about 24 hours before fixation with methanol and staining with Hema3 Manual Staining System. For each sample, ten fields of views were randomly selected and scored for the number of vacuoles.

For red/green invasion assays, coverslips with HFF monolayers were infected with 1 ml of parasites at a concentration of 2.5×10^6^ parasites/ml for 30 minutes, then were washed with DMEM three times and fixed with 4% methanol-free paraformaldehyde for 15 minutes. After being quenched with 0.1 M glycine and washed with PBS three times, the samples were incubated with mouse anti-Sag1 (1:2000) antibody in 3% BSA/PBS for 40 minutes and followed with three washes with PBS. The samples were then permeabilized and treated as described above for IFAs using rabbit anti-Mic5 (1:2000) as the primary antibody and goat anti-rabbit/mouse Alexa Fluor 594/488 (1: 2000) as the secondary antibody. For each coverslip, ten views were randomly selected for counting extracellular (green, attached but not invaded) and total (red, both extracellular and intracellular) parasites.

### Microneme secretion assay

As described before (22), freshly lysed parasites were harvested and resuspended with invasion medium (DMEM, 1.5g/L NaHCO_3_, 20mM HEPEs, pH 7.4 and 3% FBS) at a concentration of 1×10^9^ parasites/ml. Two aliquots of 100 μl of the parasite suspension were used for each sample, one for ethanol stimulated secretion and the other for natural secretion. For ethanol stimulated secretion, 2% ethanol was added to each sample. Ethanol-treated samples were incubated at 37 °C for 10 minutes, and natural secretion samples were incubated at 37 °C for 1 hour. After incubation, the samples were centrifuged at 1000 ×g for 3 minutes, and 80 μl of supernatant was collected and subjected to second centrifugation, and finally, 60 μl of the new supernatant was used for western blotting. The pellets from the first centrifugation were resuspended with PBS and centrifuged again, and the final pellets were lysed with RIPA buffer, and the lysates were used for western blotting.

### Cross-linking and immunoprecipitation

As described previously (22), freshly lysed parasites were harvested and incubated for 10 minutes in PBS with 5mM DSSO. Tris buffer was added to a final concentration of 20 mM to quench the reaction at room temperature for 5 minutes. After washes, the parasites were lysed with 1 ml RIPA lysis buffer containing a protease and phosphatase inhibitor cocktail at 4 °C for one hour. The lysate was then centrifugated, and the supernatant was incubated with 25 μl of mouse IgG magnetic beads (Cell signaling) for 1 hour at room temperature for precleaning. The unbound lysate was separated from the IgG beads and was incubated with 25 μl of anti-HA magnetic beads (Fisher Scientific) for another hour at room temperature. The anti-HA magnetic beads were washed with RIPA lysis buffer and PBS and then submitted to the Indiana University School of Medicine Proteomics Core facility for mass spectrometry.

### Intracellular magnesium concentration measurements

Freshly lysed parasites were harvested centrifugated at 1000 ×g for 10 minutes. The parasites were resuspended with 1 ml of pre-warmed Buffer A (116 mM NaCl, 5.4 mM KCl, 0.8 mM MgSO4, 5.5 mM D-glucose, and 50 mM Hepes, pH 7.4) with 5mM Magnesium Green (Invitrogen), and incubated at 37 °C for 30 min. After incubation, the parasites were washed with Buffer A two times to remove extracellular Magnesium Green, then resuspended with Buffer A at a concentration of 3×10^7^ parasites/ml and incubated at 37 °C for another 30 min to allow the complete de-esterification of intracellular AM esters. After incubation, 100 μl of parasites were loaded to each well of a 96 Well Black/Clear Bottom Plate. A Synergy Plate Reader was used to measure fluorescence with Excitation 488 nm and Emission 531 nm.

### Mouse injection and blood collection and serum antibody detection

All experiments using mice were performed with the approval of the Institutional Animal Care and Use Committee (IACUC) of Indiana University School of Medicine (IUSM) and in strict compliance with the National Institutes of Health (NIH) Guide for the Care and Use of Laboratory Animals. The 35 mice used in this study were female CBA/J mice aged 6-8 weeks, and they were divided into seven groups with five mice in each. Parasites for injection were syringe released from infected cells and resuspended in PBS to a concentration of 1000 parasites/ml or 100 parasites/ml. 100 μl of PBS with 100 or 10 parasites of each strain were injected into each mouse intraperitoneally. The control group of mice was injected with 100 μl 1xPBS.

Blood samples of surviving mice were collected from the submandibular vein. Mouse sera were used in immunoblotting assays to detect anti-*Toxoplasma* antibodies. Briefly, 5 μl of 1xPBS containing 105 freshly lysed Toxoplasma parasites were blotted onto a piece of nitrocellulose membrane and allowed to dry. The membranes were placed into 24 well plates and blocked with 5% NFDM/TBST for 45 min, then incubated with 1: 50 diluted sera isolated from the blood samples in 5% NFDM/TBST for one hour. After three washes with TBST, the membranes were incubated with anti-mouse IgG HRP in 5% NFDM/TBST for one hour. After three washes with TBST, the membranes were treated with SuperSignal West Pico chemiluminescent substrate (Pierce Chemical) and imaged using FluorChem E (Proteinsimple).

## ACKNOWLEDGMENTS

We want to thank Dr. Marc-Jan Gubbels for sharing anti-IMC3 antibodies, Dr. David Sibley and Dr. Peter Bradley for sharing plasmids, and members of the Indiana University School of Medicine Proteomics Core Facility help with the Mass Spectrometry analysis. This research was supported by the National Institute of Health grants 1R01AI123457, R01AI149766, and R21AI124067 to GA.

## SUPPLEMENTARY MATERIALS

**Supplementary Figure 1.**
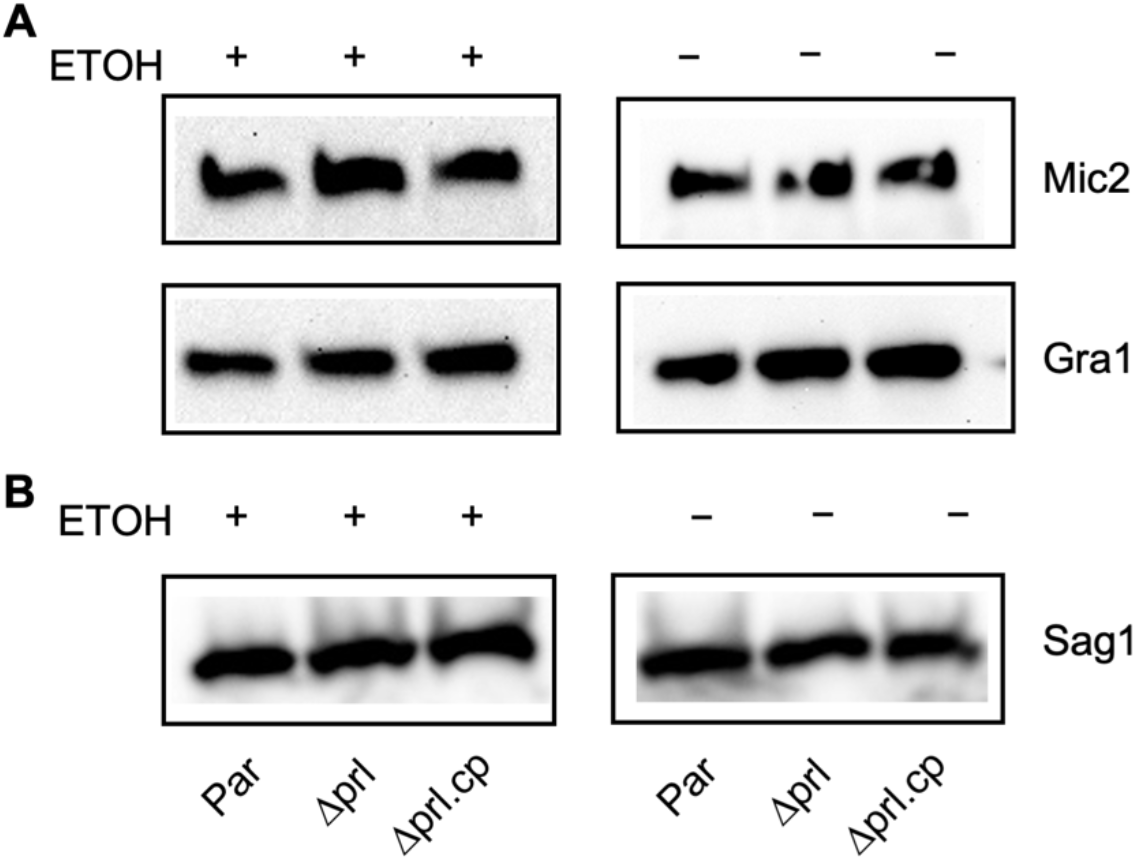
Microneme secretion in TgPRL knockout strain. Extracellular parasites of the parental (par), knockout (Δ*prl*), and complemented (Δ*prl*.cp) strains were incubated for 10 minutes with ethanol or for 1 hour without ethanol and spun down. (A) The supernatants, which contain secreted antigens, were processed for western blots with antibodies anti-TgMic2 to indicate microneme secretion and anti-TgGRA1 to indicate dense granule secretion. (B) The pellets, which contain the parasites, were lysed and processed for western blots with anti-TgSag1 to confirm an equal number of parasites were used for each treatment.

**Supplementary Figure 2.**
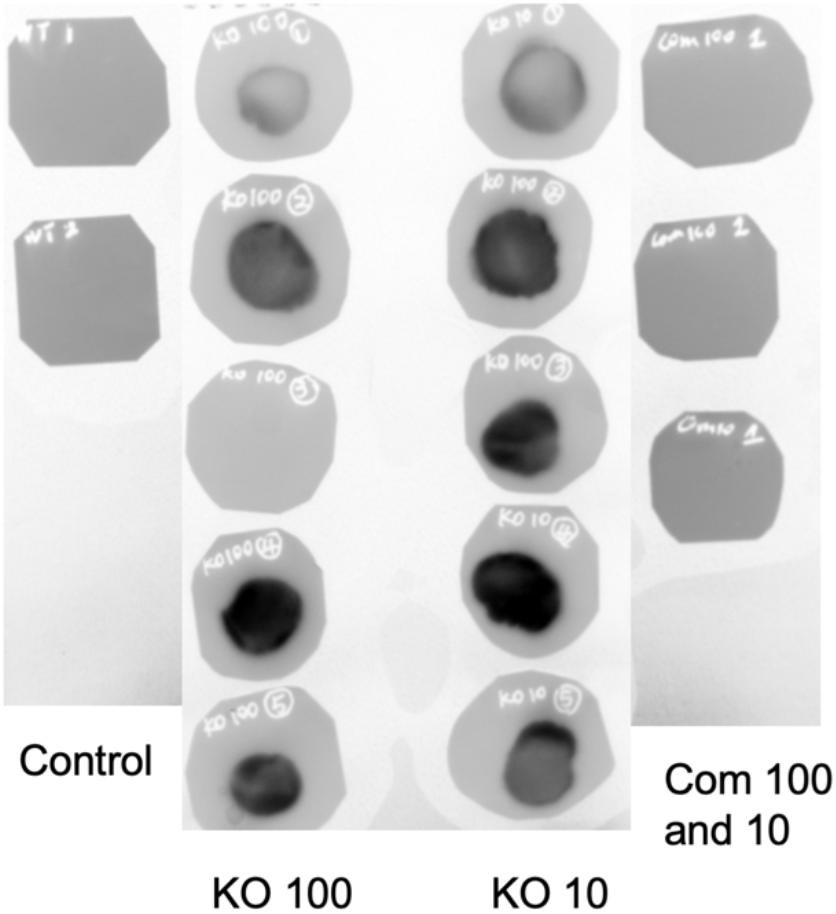
Mice infected with TgPRL knock out parasites produce anti-*Toxoplasma* antibodies. Blood was collected from the survived mice by cheek bleeding, and the serum of each mouse was used to blot with the *Toxoplasma* parasites dotted on the nitrocellulose membranes.

